# Zoomqa: Residue-Level Single-Model QA Support Vector Machine Utilizing Sequential and 3D Structural Features

**DOI:** 10.1101/2021.01.28.428680

**Authors:** Kyle Hippe, Cade Lilley, William Berkenpas, Kiyomi Kishaba, Renzhi Cao

## Abstract

**Motivation:** The Estimation of Model Accuracy problem is a cornerstone problem in the field of Bioinformatics. When predictions are made for proteins of which we do not know the native structure, we run into an issue to tell how good a tertiary structure prediction is, especially the protein binding regions, which are useful for drug discovery. Currently, most methods only evaluate the overall quality of a protein decoy, and few can work on residue level and protein complex. Here we introduce ZoomQA, a novel, single-model method for assessing the accuracy of a tertiary protein structure / complex prediction at residue level. ZoomQA differs from others by considering the change in chemical and physical features of a fragment structure (a portion of a protein within a radius *r* of the target amino acid) as the radius of contact increases. Fourteen physical and chemical properties of amino acids are used to build a comprehensive representation of every residue within a protein and grades their placement within the protein as a whole. Moreover, ZoomQA can evaluate the quality of protein complex, which is unique.

**Results:** We benchmark ZoomQA on CASP14, it outperforms other state of the art local QA methods and rivals state of the art QA methods in global prediction metrics. Our experiment shows the efficacy of these new features, and shows our method is able to match the performance of other state-of-the-art methods without the use of homology searching against database or PSSM matrix.

**Availability:** http://zoomQA.renzhitech.com

## 1 Introduction

Proteins are the drivers of biological action. They are responsible for everything from locomotion to digestion to the creation of energy. The ability for proteins to complete these functions is largely dependent on their tertiary structure: the three-dimensional arrangement of amino acids, the primary building blocks of proteins. Understanding, predicting, and analyzing protein tertiary structure is therefore a large key to breakthroughs in many different areas of biology, such as drug discovery [1, 2].

Advancements in next generation sequencing technologies allows for efficient and accurate generation of protein sequences. However, methods for determining their structure, such as X-ray crystallography and Nuclear Magnetic Resonance, are both time consuming, costly, and in some cases not possible. To address this, researchers in the bioinformatics field have developed numerous computational methods for tertiary structure prediction [3, 4, 5, 6, 7, 8, 9, 10, 11, 12]. With more and more tertiary structure predictions being made, the importance of developing methods to evaluate the quality of these models has increased. The Critical Assessment of Techniques for Protein Structure Prediction (CASP) is designed to benchmark progress of computational protein structure prediction methods and attracts hundreds of research groups around the world every other year. As CASP is gaining more exposure, private companies have begun participating in the CASP 14 competition. In 2020, companies such as Google put their resources towards protein tertiary structure prediction. Google’s latest AI algorithm, AlphaFold 2 [13] has revolutionized the field with their performance in CASP 14. Their method has the capability of making high-accuracy protein structure predictions comparable to the expensive and time-consuming lab experiments. This vast improvement in accuracy highlights the importance of evaluating the predicted protein decoy, especially the residue-level accuracy. This is referred to as Estimation of Model Accuracy problem or the protein Quality Assessment (QA) problem.

Computational methods for the QA problem aim to quantify the accuracy of a protein decoy in reference to its native structure, but without the knowledge of its ground truth. There are two metrics that address this, GDT-TS [14] (global distance test tertiary structure), which refers to the global structure accuracy, and the LDDT [15] (local distance difference test) for the accuracy of individual amino acids in the prediction. Methods for protein quality assessment are improving, as noted in the Critical Assessment of Techniques for Protein Structure Prediction (CASP) 13 experiment [16]. Current methods can be split into two general approaches [17]: single-model methods that only have access to a single prediction and must estimate its quality [18, 19, 20, 21, 22, 23] and consensus models that look at many models at a time to evaluate conserved structural patterns in order to infer quality [24, 25]. Single model methods utilize a range of different input features and methods in order to predict local or global quality. For example, DeepQA [26], a single model global method utilizing deep belief networks predicts global quality using structural, chemical, and knowledge-based energy scores to achieve good performance in the CASP 12 experiment. Similar to this, SMOQ [27] predicts absolute local qualities of tertiary structure models based on structural features (e.g., secondary structures and chemical properties (e.g., solvent accessibility). Another method, VoroMQA [28], considers a protein’s atoms and uses Voronoi tessellation of those atoms in order to predict the local quality of amino acids. The features used in most of those methods are similar and need input from both protein sequence and protein decoys, or databases for secondary structure predictions. In addition, these methods didn’t work on protein complex.

Here we propose ZoomQA, a novel single-model quality assessment method based on sequential and 3-dimensional structural and chemical features. We benchmark this tool on the CASP 14 released targets and compare its performance to other state-of-the-art methods. The accuracy achieved in local quality metrics without the use of database homology searching indicates the value of the novel features and furthers the validity of using these features to perform protein quality assessment.

The paper is organized as follows. In the Method section, we describe the data acquisition and feature generation for the tool and address the model architecture and training details. In the Results section, we analyze the performance of ZoomQA in comparison to other methods. In the Discussion section, we provide a summary and interpretation of results. In the Conclusion section, we address significant findings from this work and address future directions.

## 2 Methods

ZoomQA uses a novel representation of amino acids in protein structure and addresses the residue level protein quality assessment problem with help of machine learning techniques.

### 2.1 Data preparation

The feature input for ZoomQA is based on computational analysis of 55,000 crystal PDBs retrieved from http://www.rcsb.org/ [30] as well as CASP models (CASP6 through CASP13) from the CASP website http://predictioncenter.org/download_area/. There are over 26,000,000 residues resulting in the data coming from CASP6 through CASP13. In order to balance the representation of different qualities, we randomly shuffle all of the data into 60 batches of 430,000 training examples with a 61st batch containing the overflow. We then establish 100 bins with labels ranging from 0.0 - 0.01 all the way to 0.99-1.00 and select 600 targets for each bin, for each batch. This results in 61 smaller batches of length 60,000 that are balanced in their representation of labels. These are then used as a sample space for training data. One batch is withheld from training and used as testing data. Validation data comes from 33 targets of the CASP14 competition, representing 7,736 unique PDBs.

### 2.2 PDB Data Analysis

The basis of this work was verified by analyzing the crystal PDBs obtained from the Protein Data Bank [31]. The first area of exploration regards something we refer to as a ‘fragment structure’ which describes a portion of a protein centered around a target amino acid and includes all residues within a radius of consideration *r*, measured in angstroms, ranging from 5 to *r* angstroms. The second area of exploration is an analysis of the occurrence of torsion angle combinations for four categories: the occurrence of angles regardless of secondary structure and three values representing the occurrence of the given torsion angles when considering the secondary structure categories of alpha-helix, beta-sheet, and coil.

When analyzing the fragment structures, the first action is extracting the contact map from the PDB. Once the contact map is extracted, we need each residue’s amino acid letter code, hydrophobicity, monoisotopic mass, solvent accessibility, and isoelectric point. From this, we were able to generate what we call ‘zoom features’, a measure of a certain metric of a fragment structure as the radius of consideration increases. Examples of the relative amino acid density graphs can be found in Figure 1 [A] which demonstrate the typical environment around the amino acid Alanine and in Figure 1 [B] that shows the typical environment surrounding Tryptophan. Since we can see different trends of the amino acid environment surrounding a specific amino acid, there were trends in this data that could allow us to infer quality of an amino acid based off of not only the relative change in fragment amino acid composition, but other chemical and physical features as well. Figure 2 shows the same change of fragment structure composition over radius of consideration increase for hydrophobicity, monoisotopic mass, solvent accessibility, and isoelectric point, respectively. Once this information is acquired, we can compile the data for the change over radius expansion. If we consider amino acid at index 0 in the sequence, most methods consider the sliding window of neighbors to that amino acid in the sequence (i.e the residue at index 1, 2, 3, etc.) [32]. However, since proteins are not strictly linear, other amino acids not directly next to the target amino acid in sequence play a role in determining the target amino acid’s placement. More importantly, it will be difficult for protein complex where several chains are not directly connected. To address this, we analyze the 3D radius around an amino acid as the target’s environment. We can extract relevant data from this environment which is a more representative feature of how amino acids are placed within proteins.

**Figure 1:**
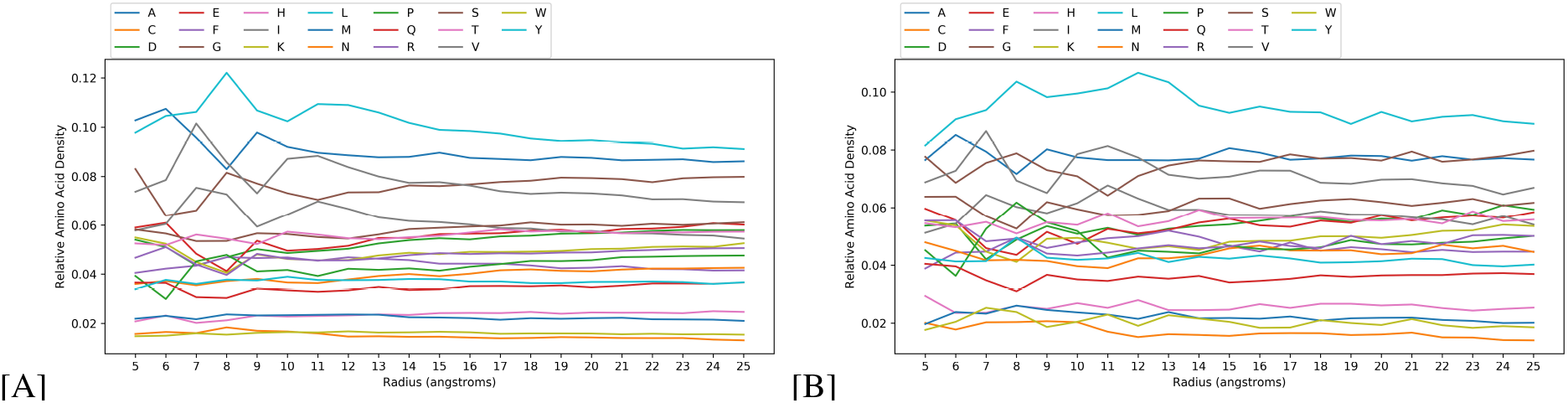
(A) The relative density of amino acids around a target amino acid (Alanine) as the radius of consideration increases from 5 to *r* (B) The relative density of amino acids around a target amino acid (Tryptophan) as the radius of consideration increases from 5 to *r*

**Figure 2:**
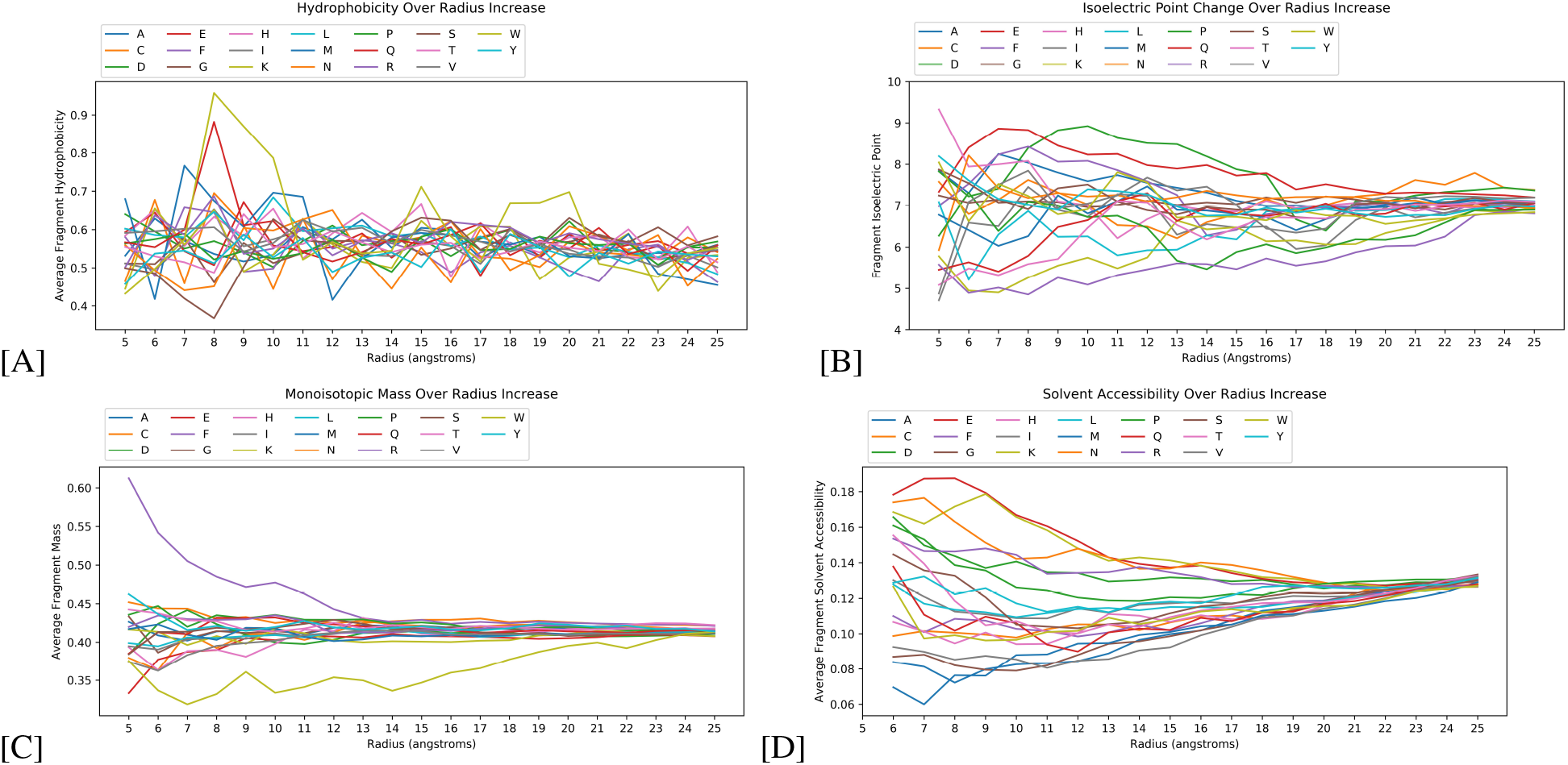
(A) The average hydrophobicity of the fragment structure at radius 5 to *r* centered around the target amino acid with letter codes of amino acids represented in the figure legend. (B) The average monoisotopic mass of the fragment structure at radius 5 to *r* centered around the target amino acid with letter codes of amino acids represented in the figure legend. (C) The average solvent accessibility of the fragment structure at radius 5 to *r* centered around the target amino acid with letter codes represented in the figure legend. (D) The fragment structure at radius 5 to *r* centered around the target amino acid with letter codes represented in the figure legend. This is generated by the IsoelectricPoint module in Python’s package BioPython [29].

The second area of exploration was an analysis of dihedral angles that occur in crystal PDBs. In theory, both phi and psi can be in a range (−180, 180). However, in practice, the torsion angles cannot reach that full range. This is due to the amine group of each amino acid. The size, mass, hydrophobicity, and other factors all play into the placement of the amine group, and the placement of the amine group can sterically prohibit certain angles from existing in nature. Furthermore, torsion angles correlate with the secondary structure of the amino acid and can be used to infer secondary structure. This feature is often used in quality assessment tools like DeepQA [26], AngularQA [33], and SMOQ [27], but is oftentimes encoded as a 0 or 1 describing whether or not the torsion angles represent the predicted secondary structure. We can get a more representative feature of torsion angles and secondary structure by analyzing the occurrence of these torsion angles regardless of secondary structure and the torsion angle occurrences when the amino acid is within secondary structures (alpha-helices, beta-sheets, and coils). Figure 3 and Figure 4 show the Ramachandran plots that represent the stability scores for two different amino acids: alanine and proline. Both amino acids exhibit different common torsion angles dependent on the secondary structure they are in. These different values led us to believe that these different values could help our model differentiate the quality of an amino acid based off of its predicted secondary structure and its predicted stability within that structure.

**Figure 3:**
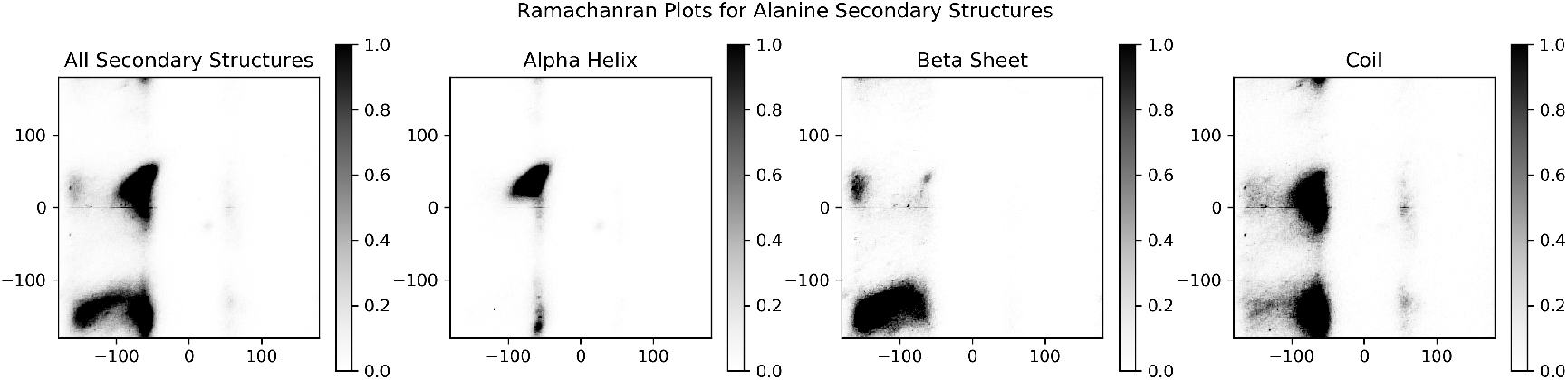
Ramachandran Plot for Alanine. These plots represent the torsion angles found *in vivo* for Alanine from left to right, regardless of secondary structure is an alpha helix, a beta sheet, or a coil.

**Figure 4:**
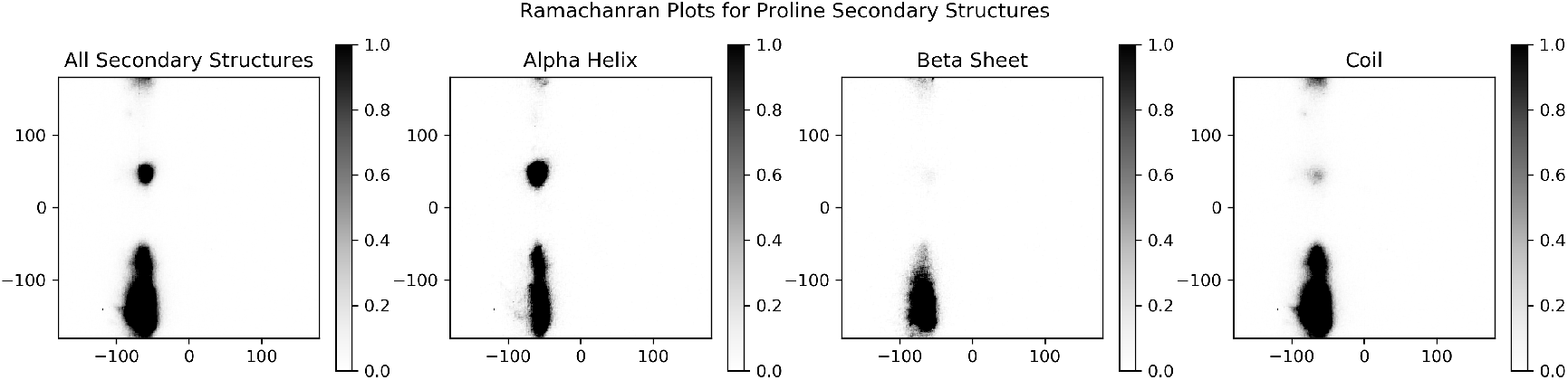
Ramachandran Plot for Proline. These plots represent the torsion angles found *in vivo* for Proline from left to right, regardless of secondary structure is an alpha helix, a beta sheet, or a coil.

In order to generate this stability score, we explored various machine learning techniques before settling on Random Forest regressors. For each amino acid, a Random Forest was trained on 97200 instances of angles with their normalized relative occurrence and tested on the remaining 32400 instances of angles that can come from the combination of angles in the range (−180, 180). After exploring hyperparameters of the number of trees and maximum tree depth, the optimal parameters were found by minimizing the testing average error + maximum error + the difference between the testing and training values for maximum and average error.

### 2.3 Features

ZoomQA utilizes eighteen features regarding the chemical and physical properties of the target amino acid and its environment. To reduce confusion, we will use the term ‘fragment’ to describe a region of a protein that can be generated by including all amino acids within a radius of consideration *r* of a target amino acid where *r* represents a distance measured in angstroms. Two data sets were generated, one where *r* was set to 25 angstroms, and another where *r* was set to 55 angstroms. Unless the description states otherwise, all described features are normalized by the equation:

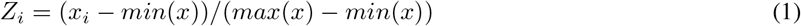

Where *Z_i_* is the normalized value, *x_i_* is the value we are trying to normalize, *min*(*x*) represents the minimum value from the set *x*, and *max*(*x*) represents the maximum value from the set *x*.

The first feature is the average amino acid density of a fragment as the radius of consideration increases from 5 angstroms to *r* angstroms. This is a *r* x 20 matrix. The columns of this matrix correspond to the letter codes for all twenty amino acids in alphabetical order. The rows of this matrix represent the radius of consideration in the set 5-*r* (e.g row 0 represents a radius of consideration of 5 angstroms, row *r* represents a radius of consideration of *r* angstroms). Each element of this matrix represents the relative density of the column amino acid in the fragment with a radius equal to the row + 5 angstroms from the center amino acid.

The second feature is the average hydrophobicity of the fragment of protein in contact with the target amino acid as the radius of consideration increases from 5 to *r* angstroms. This is a vector with length *r*, where index 0 represents the average hydrophobicity of all amino acids within the fragment if the radius of consideration is 5 angstroms. This includes the hydrophobicity of the target amino acid, as that influences the overall hydrophobicity of the structure.

The third feature is the average monoisotopic mass of the fragment of protein in contact with the target amino acid as the radius of consideration increases from 5 to *r* angstroms. This is a vector with length *r*, where index 0 represents the average mass of all amino acids within the fragment if the radius of consideration is 5 angstroms. This includes the mass of the target amino acid because it influences the overall mass of the structure.

The fourth feature is the average solvent accessibility of the fragment of protein in contact with the target amino acid as the radius of consideration increases from 5 to *r* angstroms. This is a vector with length *r*, where index 0 represents the average solvent accessibility of all amino acids within the fragment if the radius of consideration is 5 angstroms. This includes the solvent accessibility of the target amino acid as that influences the overall solvent accessibility of the structure.

The fifth feature is the isoelectric point of the fragment of protein in contact with the target amino acid as the radius of consideration increases from 5 to *r* angstroms. This is a vector with length *r*, where index 0 represents the isoelectric point of the fragment generated by a radius of consideration of 5 angstroms. This is generated by the IsoelectricPoint module in Python’s package BioPython [29].

The sixth feature is a length *r* vector that represents the average distance of all amino acids to the target amino acid as the radius of consideration increases from 5 to *r*. This value is normalized by the radius of gyration, defined as the maximal distance between the target amino acid and any amino acid within the radius of consideration. This is accompanied by the seventh feature which is a length *r* vector that represents the standard deviation in the distance between the target acid and its set of amino acids in contact as the radius of consideration increases. The input data is normalized before calculating the standard deviation, thus the feature is normalized upon creation. The input data in normalized by dividing all distances between the target amino acid and its contacts by the radius of gyration.

The eighth feature is a vector with length *r* that represents the percentage of the protein as a whole that is within the radius of consideration as the radius increases from 5 angstroms to *r* angstroms.

The ninth feature is very similar to the first feature where we find the relative density of each amino acid for fragments as the radius of consideration increases. For this feature, we weight the occurrence of the amino acids, adding an increased weight if the amino acid in contact is not sequentially in contact with the target amino acid. This generates a contact matrix that emphasizes the contacts resulting from the folding of a protein.

The tenth feature is the stability score of the target amino acid’s torsion angles as generated by the random forest models described in the previous section. We generate four stability scores. The first score is the stability score of the angles regardless of the secondary structure. The next three scores are the stability scores of the torsion angles if the secondary structure of this amino acid was an alpha-helix, beta-sheet, or coil, respectively. We do not consider the secondary structure of this amino acid coming from the PDB, however.

The final features all pertain to the center amino acid, the target. We include the amino acids’ monoisotopic mass, the hydrophobicity, the solvent accessibility, isoelectric point, and the torsion angles (two values), all normalized to values between 0 and 1. We also include the amino acid letter code and the secondary structure extracted from protein structure as one-hot encoded vectors.

### 2.4 Method Architecture and Prediction Process

The final model was trained on 60,000 examples of data generated from a maximum radius of consideration of 55 angstroms. This led to a total of 2,397 features being generated for each amino acid in the training set. The final model selects the top 100 performing features based on their Pearson correlation to their labels. The Support Vector Machine is trained as the final model with the RBF kernel, a C value of 1.0, an epsilon value of 0.1, and a gamma value of 1.0.

Figure 5 represents the overall process of creating local quality predictions based off of a single PDB input. In order to use ZoomQA, PDB format input is required (the input could be single chain protein or complex). From there, each PDB is loaded and features are extracted and compiled into a 47×51 dimension matrix. Once this is created, we select the top 100 correlated features as we found from our experiments. Then, each amino acid is fed through the support vector machine producing a local quality score. This is repeated for all amino acids in the structure. Once all local quality scores are calculated, the GDT-TS score [34, 14] is calculated using the equation:

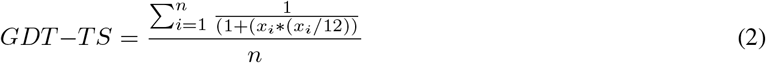

**Figure 5:**
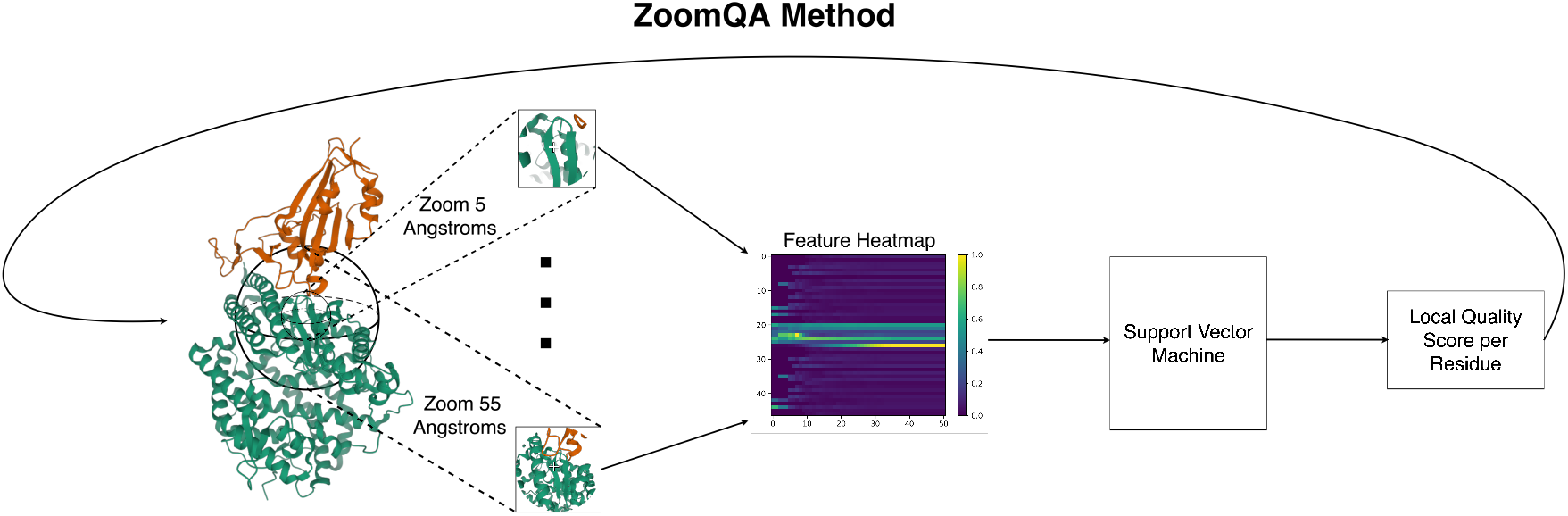
Flowchart of ZoomQA method as a whole. PDB data is taken in and features are generated. Each feature set is then fed into a support vector machine that predicts the local quality of that amino acid. Repeat this process for all amino acids in the PDB.

Where the set *x* is set of local distance error predictions for each amino acid.

## 3 Results

ZoomQA was benchmarked on the latest CASP 14 dataset and compared with other top-performing single model QA methods. In total, 33 targets are used from CASP 14 dataset, and all predictions are filtered so they have the same sequence in each target. Since ZoomQA can predict on both global and local quality, we evaluate GDT-TS using standard metrics of per-target average Pearson correlation, per-target Kendall Tau correlation, per-target Spearman correlation, per-target loss, overall Pearson correlation, overall Spearman correlation, and overall Kendall Tau correlation. For local quality metrics, we evaluate the minimum, average, and maximum GDT-TS error and LDDT error, as well as the GDT-TS and LDDT standard deviation in error for targets over a distribution of bins.

Figure 6 illustrates the performance of ZoomQA compared to other methods’ performances on global metrics in the overall CASP 14 benchmark set. Figure 7 represents the performance of the stage 2 (top 150 predicted models are selected for each target in stage 2) CASP14 benchmark set on global metrics. As we can see from Figure 7, ZoomQA betters the performance of SMOQ on all per-target correlation metrics, as well as performing better in the per target loss, and nearly matches the performance of DeepQA in the loss metric.

**Figure 6:**
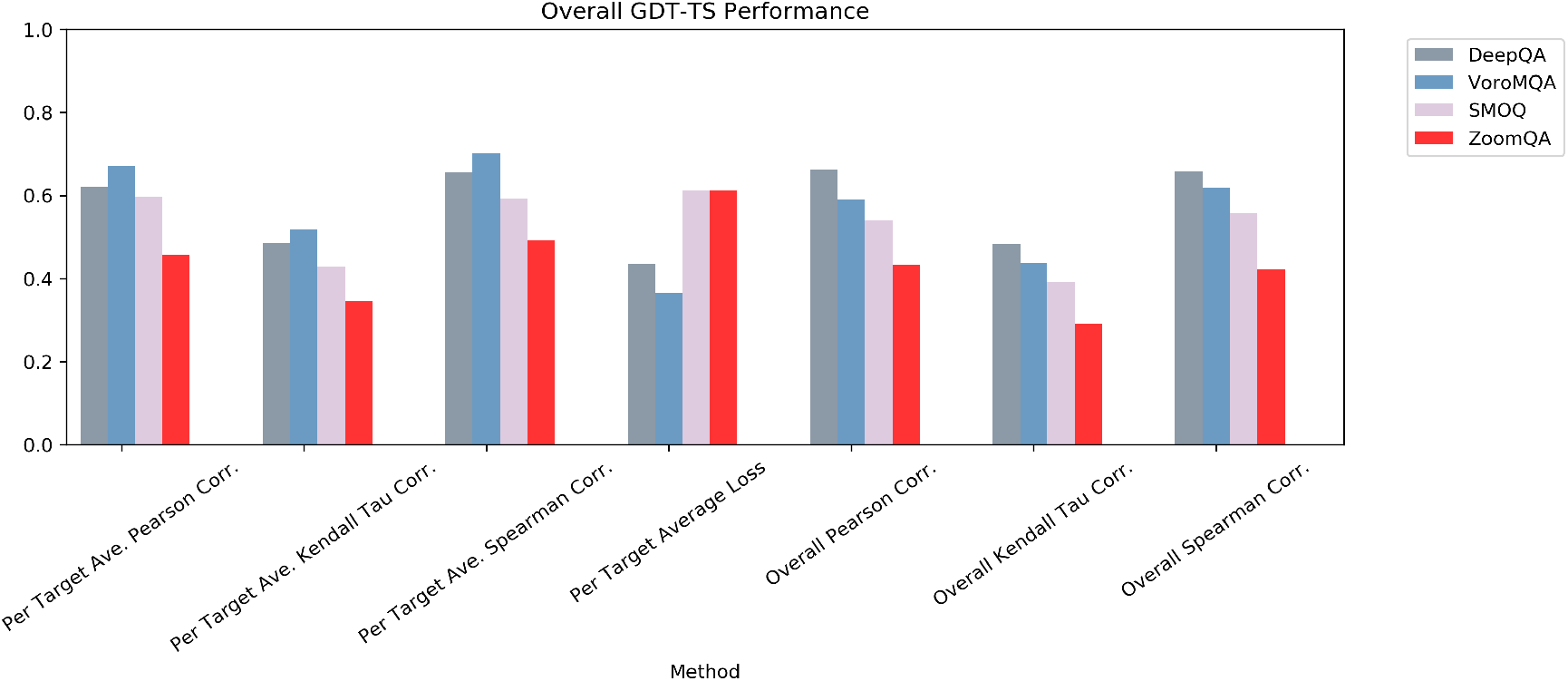
The overall GDT-TS performance of various methods tested on the CASP 14 benchmark dataset.

**Figure 7:**
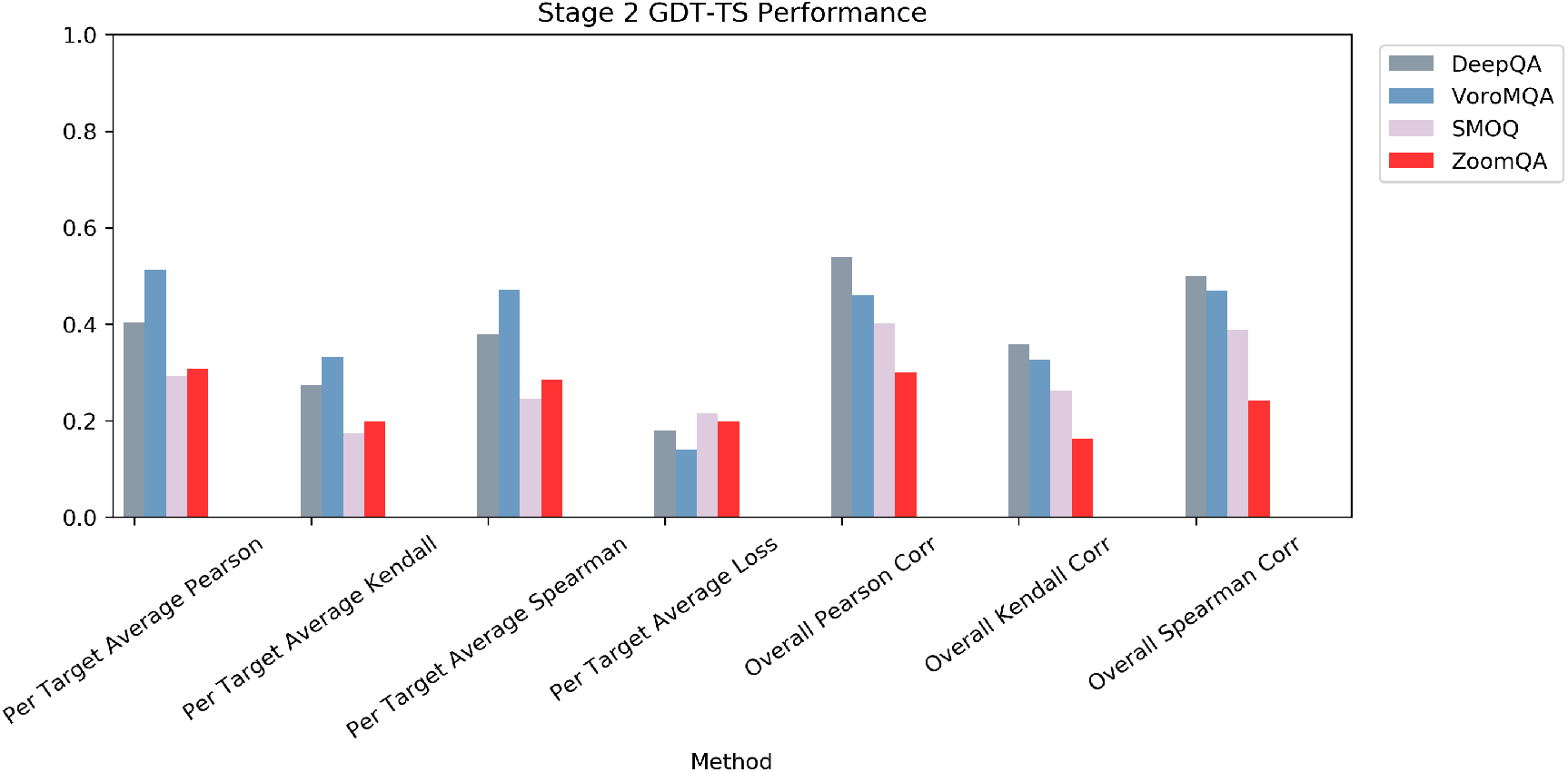
The Stage 2 GDT-TS performance of various methods tested on the CASP 14 benchmark dataset.

Moving onto performance in local quality, Table 3 demonstrates minimum, average, and maximum distance error for the other local QA methods tested, while Table 3 represents the minimum, average, and maximum LDDT error for the overall CASP 14 benchmark set. When measuring distance error, ZoomQA falls short of other methods but manages to be within ten percent of each method for minimum, average, and maximum error. When looking at overall LDDT performance, however, ZoomQA manages to beat both VoroMQA and SMOQ in all metrics. As seen by Figure 9, ZoomQA outperforms both SMOQ and VoroMQA for all LDDT metrics. This trend continues for Stage 2 performance. ZoomQA is beaten in Stage 1 performance only by the method SMOQ on the average and maximum LDDT error, but still outperforms VoroMQA here. In the GDT-TS metric, ZoomQA nearly matches performance with SMOQ and VoroMQA for stage 1 predictions. Figure 8 demonstrates the standard deviation of distance error when the target value is in the range 0 to 19 angstroms. ZoomQA outperforms SMOQ and VoroMQA at ranges 1 to 7. It is worth noting that all distances greater than 19 angstroms were grouped into the final bin, causing a large spike in the deviation of distance error. Figure 9 shows the standard deviation of LDDT error when the target value is in 20 bins, 0-0.05, 0.05-0.010 and so on. ZoomQA outperforms DeepQA for all bins and outperforms SMOQ at bins 0-14.

**Figure 8:**
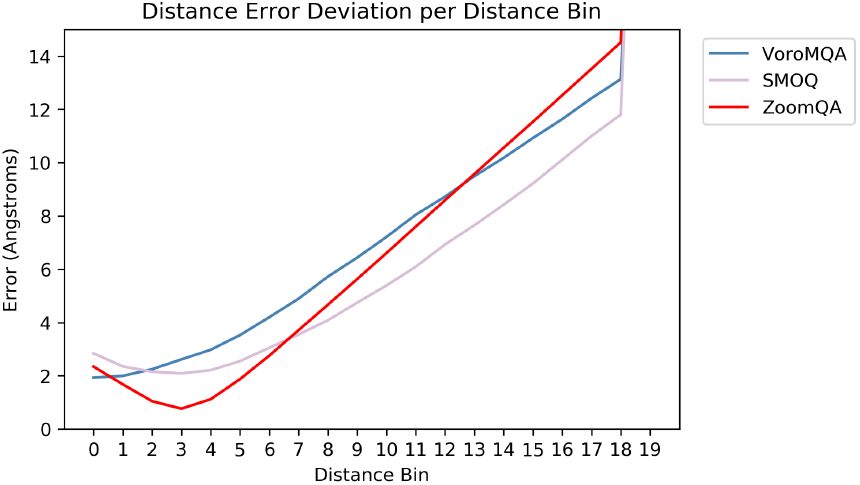
Performance of ZoomQA vs other local QA methods. Each line represents the standard deviation of the absolute distance error when the ground truth value is in each bin.

**Figure 9:**
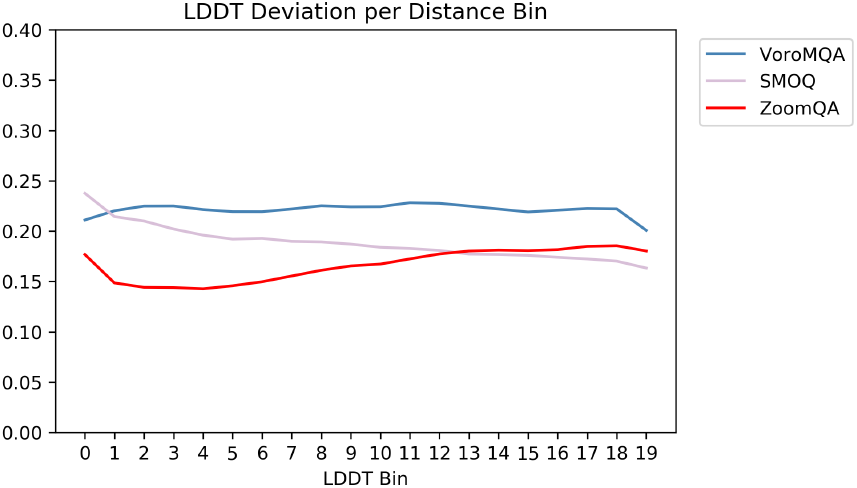
Performance of ZoomQA vs other local QA methods. Each line represents the standard deviation of the LDDT error when the ground truth value is in each bin.

## 4 Discussion

### 4.1 Feature Selection and Training

With the generation of all of the features described in Section 2.3, there are a number of data points which are not correlated to the outputs and impede the convergence of a model regardless of the architecture. To get around this, we selected features utilized in the model based off of their Pearson correlation and selected features based off of Pearson correlation and feature clusters.

| Local QA Absolute Distance Error of ZoomQA vs other tools. |  |  |  |  |
| --- | --- | --- | --- | --- |
| Method | Stage | Minimum Error | Average Error | Maximum Error |
| ZoomQA | Overall | 1.156 | 12.668 | 248.751 |
|  | Stage 1 | 1.466 | 22.32 | 248.751 |
|  | Stage 2 | 1.156 | 8.53 | 111.494 |
| SMOQ | Overall | 1.353 | 12.016 | 242.104 |
|  | Stage 1 | 1.551 | 20.375 | 242.104 |
|  | Stage 2 | 1.353 | 8.307 | 104.962 |
| VoroMQA | Overall | 1.094 | 11.798 | 234.13 |
|  | Stage 1 | 1.381 | 19.844 | 234.13 |
|  | Stage 2 | 1.094 | 8.290 | 100.98 |

| Local QA LDDT Error of ZoomQA vs other tools. |  |  |  |  |
| --- | --- | --- | --- | --- |
| Method | Stage | Minimum Error | Average Error | Maximum Error |
| ZoomQA | Overall | 0.063 | 0.159 | 0.61 |
|  | Stage 1 | 0.069 | 0.18 | 0.43 |
|  | Stage 2 | 0.0635 | 0.151 | 0.402 |
| SMOQ | Overall | 0.064 | 0.189 | 0.718 |
|  | Stage 1 | 0.075 | 0.159 | 0.355 |
|  | Stage 2 | 0.095 | 0.199 | 0.415 |
| VoroMQA | Overall | 0.079 | 0.218 | 0.824 |
|  | Stage 1 | 0.082 | 0.12 | 0.435 |
|  | Stage 2 | 0.099 | 0.228 | 0.546 |

#### 4.1.1 Method 1: Pearson Correlation

When selecting the features based on feature correlation, we calculate Pearson correlation on each feature and select the top *n* features based on the absolute value of their feature correlation. Once the features are selected, the features are arranged either into a vector with the top correlated feature at the beginning of the vector and the lowest at the end, or into an nxm matrix where coordinate (0,0) represents the highest correlated feature and the lowest correlation is at coordinate (n,m). The distinction between creating a vector or a matrix depends on the method being tested. Vectors are created when training random forests, support vector machines, and multi-layer perceptrons, whereas matrices are used when training convolutional neural networks. This method was used in the final model and achieved all the metrics stated in Section 3.

#### 4.1.2 Method 2: Pearson Correlation and Clustering

When selecting features based on correlation and clustering, we repeat the process described above and rank each feature based on their Pearson correlation. Next, we perform K-Mean clustering on each feature and generate n number of clusters of features, while n is the number of features you want to select. In our case, n is 100 since we would select top 100 features. Each feature is selected from each cluster prioritizing features near the centroid until we have n features. These features are then arranged into a vector with the highest correlation feature in the first index and the lowest correlation feature in the last index, or into an *n*x*m* matrix where coordinate (0,0) represents the highest correlated feature and the lowest correlation is at coordinate (n,m). The distinction between creating a vector or a matrix depends on the method being tested. Vectors are created when training random forests, support vector machines, and multi-layer perceptrons, whereas matrices are used when training convolutional neural networks. Ultimately, selecting features based on Pearson correlation and clustering resulted in very few effective features being chosen. When the clusters were generated, a select few of the clusters held the majority of the high correlation features. When choosing features from each cluster, we very quickly exhausted the pool of effective features. Using this technique for feature selection resulted in an average absolute distance error of 14.03 angstroms, slightly worse performing than selecting solely based on Pearson correlation.

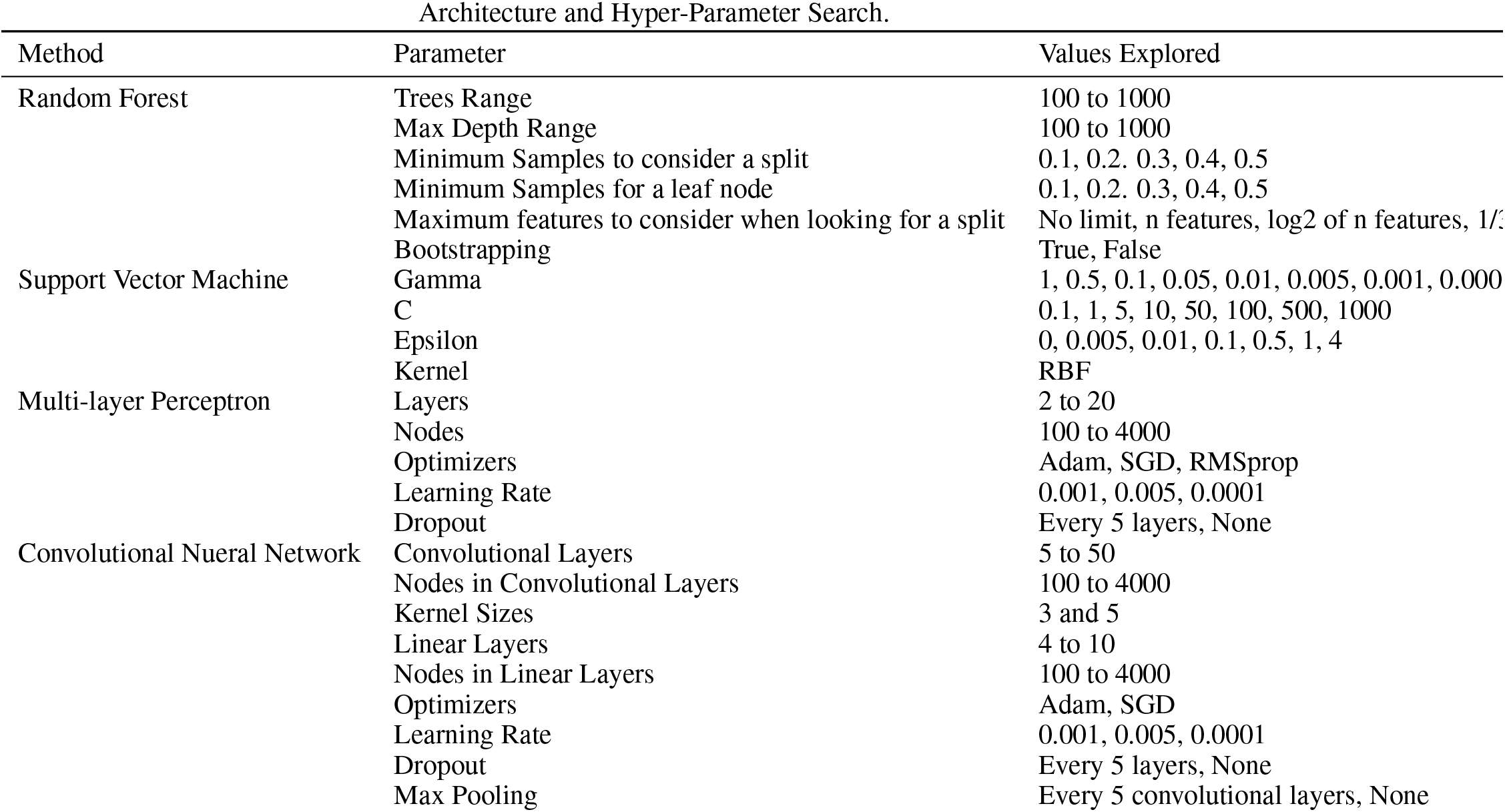

#### 4.1.3 Architecture and Hyper-Parameter Search

Once the highly correlated features were chosen, extensive architecture and hyper-parameter searches were conducted. Table 4.1.3 describes the machine learning techniques and the hyper parameters explored. This extensive search was conducted only on data generated with a maximum radius of consideration of 25. This extensive of testing was not possible on the data set generated with a maximum radius of consideration of 55, which ended up being the final model due to time and resource constraints.

#### 4.1.4 Efficacy of Features

When using all of the features for training, we achieved a mean absolute distance error of 16.23 angstroms. During efforts to improve these results, we found a large number of features that exhibited very low Pearson correlation with their labels. Upon this realization, we began filtering our feature set based on the features’ personal correlation with the labels. We would like to demonstrate the efficacy of our ZoomQA features. When evaluating all features, there are 28 features that have over a 0.20 correlation with the targets. Once we go down to the top 100 features, this correlation only drops to 0.12. Of the top 100 correlated features, 51 come from the change in solvent accessibility as the radius of consideration increases, 28 come from the average distance of amino acids in a fragment to their center, 13 come from the change in hydrophobicity of protein fragments as the radius of consideration increases, all of the secondary structure stability scores are included, as are the hydrophobicity, mass, and isoelectric point values for the target amino acid. It is interesting to note that while the isoelectric point values for the target amino acid are included, data regarding the change in isoelectric point of the fragment structures around the target amino acid is not included. In fact, the change in isoelectric point of protein fragments ranks around the 1700 most influential features with an average correlation of around 0.02. This indicates that this feature is not indicative of the quality of an amino acid and was part of the initial troubles when training a model. This is an interesting conclusion since all other zoom features were at a minimum in the top third of features based off their correlation. Other features that rank below 0.05 correlation include certain values coming from the change in amino acid density of protein fragments and portions of the structure contact matrix. This was expected as a considerable number of targets do not contain certain amino acids, meaning there are large portions of the training set that contain all zeros for whole rows of data. The usage of these features improved our mean absolute distance error by roughly 4 angstroms to a value of 12.67 angstroms of error.

#### 4.1.5 Performance

In regards to performance, ZoomQA manages to outperform other well-performing models on the CASP 14 benchmark dataset on the LDDT metric and rivals their performance on the local distance metric. This performance is further highlighted in Figure 8 where ZoomQA achieves lower distance error deviation across many of the true values below 7 angstroms and in Figure 9 where it outperforms other methods in LDDT deviation when below 0.60. This is done without the use of a PSSM and any alignment data obtained from BLAST/PSI-BLAST. This highlights the efficacy of the features obtained from ZoomQA and also allows for the tool to be easier to use than other methods that would require the use of a large database or the time consuming proteins of performing sequence alignment. It also takes out some of the variation in results as sequences without homology in the provided database could harm the performance of methods that require them. ZoomQA does not perform as well as other methods in the GDT-TS calculations, but given that it manages to match the performance on stage 2 data with SMOQ (a local QA method) by only using a linear transformation to calculate the GDT-TS, this result is acceptable and can be improved upon in further iterations of the tool by including a predictor for GDT-TS.

## 5 Conclusion

In this paper, we purpose a new residue-level protein model quality assessment tool, ZoomQA, which utilizes novel sequential and 3-dimensional structural features to grade the local quality of a tertiary structure prediction. It outperforms state-of-the-art methods in local quality, particularly when measuring LDDT and rivals the performance of these methods in GDT-TS metric. Our method only needs protein structure as input, and it points out a new direction for evaluating the quality of predicted protein decoy and also protein complex.

In the future, we plan on fine-tuning the zoom features to minimize the production of low correlated features. This would eliminate the need for feature selection based on Pearson correlation. Additionally, we plan on incorporating more zoom features to better describe the 3-dimensional structure of proteins. We would also like to explore different forms of deep learning to gain insights into our data that may not be possible with conventional machine learning techniques.

## Acknowledgements

This work is supported by the Natural Sciences Undergraduate Research Program at Pacific Lutheran University.

